# *De novo* design of a safe and potent respiratory syncytial virus immuno-focusing antigen

**DOI:** 10.64898/2026.01.28.702448

**Authors:** Woo Yeon Hwang, Jinung Song, Jiwon Choe, Keun Bon Ku, Hyung-Sun Kim, Gun Young Yoon, Do Yeon Kim, Mi-Ra Choi, Eun-Ji Kim, Jae Seung Lee, Sungjun Park, Sung Kyun Lee, Seung jun Kim, Bonsu Ku, Dae-Gyun Ahn, Kyun-Do Kim, Chonsaeng Kim, Han Na Suh, Juyong Lee, Ho-Chul Shin, Junsu Ko, Young-Chan Kwon

## Abstract

Respiratory syncytial virus (RSV) remains the leading cause of severe respiratory infections in infants, the elderly, and the immunocompromised. Although stabilized full-length pre-fusion (pre-F) protein vaccines are promising, enhanced respiratory disease (ERD) remains a critical safety concern. Here, we used artificial intelligence to design a *de novo* immuno-focused antigen that structurally preserves the RSV F head region containing critical neutralising epitopes— site Ø, II and V–while replacing the non-neutralising stem with a computationally designed scaffold to minimise immunopathological risk. The lead candidate, aRF6, elicited robust protective immunity against RSV in mice and similar immunogenicity in non-human primates without detectable toxicity. Importantly, in stringent ERD-promoting models, aRF6 induced minimal pulmonary pathology and markedly attenuated Th2-biased cytokine responses, outperforming formalin-inactivated RSV and full-length-stabilized pre-F. The results of cryoelectron microscopy confirmed that the aRF6 structure precisely matched the computational predictions. These results demonstrated that computationally designed *de novo* immuno-focused antigens can yield safe and effective RSV vaccines, thereby providing a rational framework for next-generation vaccine development.

## Introduction

Respiratory syncytial virus (RSV) is the leading cause of severe lower respiratory tract infections in infants, the elderly, and the immunocompromised, resulting in an estimated 100,000 in-hospital deaths annually worldwide ^1,2^. Natural infection induces only short-lived immunity, underscoring the urgent need for a safe and durable vaccine ^3,4^. However, RSV vaccine development has been hampered for decades by a major safety setback: a formalin-inactivated RSV (FI-RSV) vaccine tested in the 1960s caused vaccine-enhanced respiratory disease (ERD) upon subsequent RSV infection ^5^. This adverse outcome is characterised by a skewed Th2-type immune response and generation of poorly neutralising antibodies (nAbs), leading to severe illness and fatalities in vaccinated children ^6,7^.

Recent efforts to develop vaccines have focused on the viral fusion (F) glycoprotein, which facilitates viral entry and is conserved across RSV subgroups ^8^. Although the pre-fusion (pre-F) conformation contains the most potent neutralising epitope, site Ø, which is the target of the recently licensed monoclonal antibody (nirsevimab), it is inherently unstable and prone to irreversible structural transitions to the post-fusion (post-F) form that obscures key antigenic sites (Fig. 1a) ^9,10^. The structural stabilization of pre-F has enabled the development of licensed vaccines for adults, the elderly, and pregnant women^11-14^. However, recent clinical trials of mRNA-based pre-F vaccines using stabilization approaches have revealed safety issues in RSV-naïve infants, with higher severe RSV-associated lower respiratory tract infection–related hospitalizations observed in vaccine recipients than in placebo recipients ^15^. This suggests that current stabilization strategies are not sufficient to fully mitigate immunopathological risk in the most vulnerable age group.

**Fig. 1.**
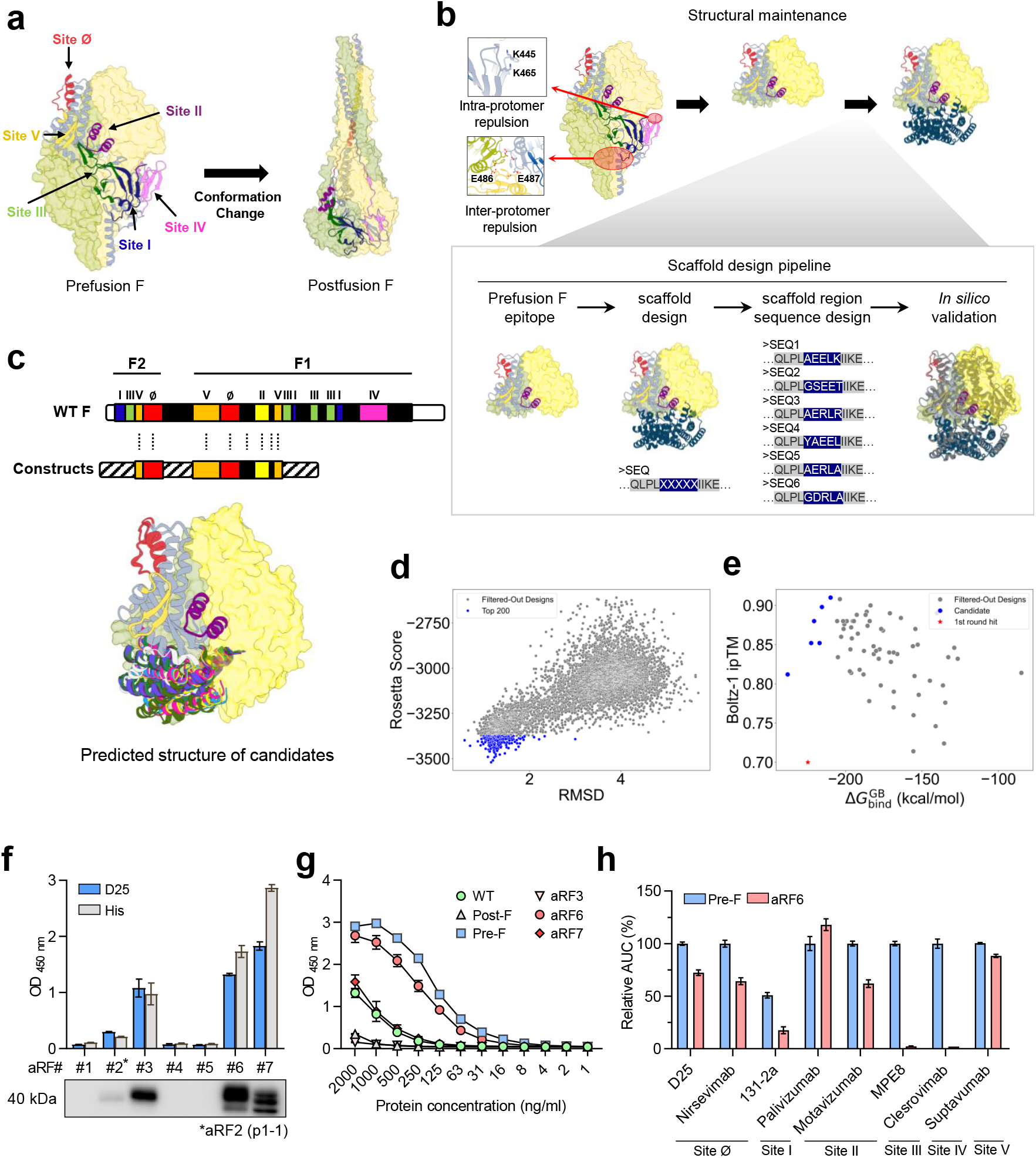
Structural de novo design of RSV F vaccine antigens and characterisation of candidate construct expression and antigenicity. **a**, Crystal structure of the pre-fusion (pre-F; PDB:4JHW) and post-fusion (post-F; PDB:3RRR) states of the RSV F glycoprotein. Known epitopes are coloured and labelled. **b**, Overall workflow of the protein design pipeline. **c**, Sequence diagram of the native RSV F protein and *de novo* antigens and aligned AlphaFold-Multimer structures of the final experimental candidates. **d**, Stability and energy metrics of candidates resulting from the expanded design phase. Scatter plot of RMSD values between designed and predicted structures and Rosetta energy scores of predicted structures. The selected 200 designs are shown in blue. **e**, Scatter plot of interfacial binding affinity and Boltz-1 ipTM of top candidates. The candidates selected for experimental validation are shown in blue. **f**, The designed F antigen constructs (aRF1–aRF7; aRF2 corresponds to p1-1) were expressed in CHO cells, and the expression levels in the culture supernatant were quantified using ELISA with anti-His and D25 antibodies. Detection was confirmed using immunoblotting. **g**, After column purification of the selected F antigen constructs (aRF3, aRF6, and aRF7), site Ø expression was assessed using a D25 antibody in a concentration-dependent manner. Wild type (WT) RSV F, post-F, and pre-F strains were used as controls. **h**, Antigenic profiling of aRF6 and pre-F was performed using a panel of site-specific monoclonal antibodies. Relative area under the curve values (%) were determined using ELISA for antibodies targeting RSV F antigenic sites Ø (D25, nirsevimab), I (131-2a), II (palivizumab, motavizumab), III (MPE8), IV (clesrovimab), and V (suptavumab). Stabilized pre-F was set to 100% for normalisation for all except 131-2a, which was normalised to post-F. The experiments related to (e)–(g) were repeated twice, and representative data are shown and presented as mean ± SE.

To overcome these limitations, we engineered novel immuno-focused RSV F protein antigens using an artificial intelligence (AI)-based *de novo* protein design strategy integrated with high-accuracy structural prediction. Our strategy was guided by three principal objectives: (1) to stabilize the trimeric RSV pre-F head while preserving its critical neutralising epitope, site Ø, together with functionally important sites II and V, which collectively dominate the neutralising antibody response ^16-19^; (2) to minimise undesired non-neutralising epitopes by selectively removing the stem region, which might contribute to aberrant immunity or ERD-like pathology; and (3) to incorporate a *de novo*–designed protein scaffold that maintains integrity and optimal presentation of these neutralising sites while minimising unintended antigenic targets.

By leveraging AI-driven protein design methods, we developed a stable immuno-focused RSV F protein antigen that incorporates a novel scaffold engineered to promote a potent immune response. We validated the structural integrity of the design using *in vitro* assays and evaluated its efficacy and safety in mouse and non-human primate (NHP) models. The AI-designed antigen provided protection comparable to that of the stabilized full-length pre-F, while compared to FI-RSV and stabilized full-length pre-F, it significantly reduced the expression of ERD biomarkers. It elicited high neutralising antibody (nAb) titres without inducing aberrant or redundant immune responses and produced potent sustained neutralisation in NHPs without observable toxicological effects. Collectively, these findings demonstrated that *de novo* antigen design can overcome both structural instability and immunological risks that have long hindered RSV vaccine development, offering a promising pathway toward safe, effective, and broadly protective immunisation for vulnerable populations worldwide.

## Results

### AI-guided *de novo* design and selection of an immuno-focused RSV F antigen

The RSV F glycoprotein mediates viral entry into host cells. Site Ø, a pre-F-specific epitope, is the most promising neutralising target; however, it becomes progressively masked during the irreversible transition to the post-F conformation (Fig. 1a). Therefore, our *de novo* design strategy focused on preserving this critical epitope alongside the functionally competent sites II and V that elicit potent nAbs. This immuno-focusing strategy comprises four steps: preservation of the epitope conformation, rational redesign of the stem region scaffold, sequence generation of the scaffold, and *in silico* validation (Fig. 1b). Adopting this approach, we first retained the potent antigenic epitopes, sites Ø, II, and V, by incorporating the pre-F conformation of the RSV F protein, while truncating the stem region to eliminate known inter-and intra-protomer repulsions ^11,20-22^. Next, we scaffolded these epitopes to generate stable immuno-focused antigens using RF diffusion ^23^ and ProteinMPNN ^24^. Subsequent *in silico* analysis demonstrated that the *de novo-*designed F proteins, which comprise 69.9% of the wild-type F protein length and sharing only 44.6% sequence identity, stably maintain the conformations of antigenic sites Ø, II, and V within their trimeric structures (Fig. 1c).

*In silico* screening was performed in two sequential phases to progressively prioritize the most promising candidates. In phase 1, candidates were ranked by their structural fidelity to the initial designs and stability of the predicted trimer structures. The latter was estimated by the interfacial binding free energy 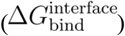 between one monomer and the remaining dimer (Extended Data Fig. 1). A detailed description of our pipeline design is presented in the Methods. From this initial phase, five antigens with the lowest 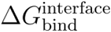 and accessible termini were selected for *in vitro* screening (Extended Data Fig 2). Four antigens were detected using anti-His immunoblotting; however, their expression levels were insufficient for further characterisation. Despite this, p1-1, which had the lowest 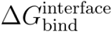 among the five, exhibited the highest binding to the D25 antibody, recognizing the antigenic site Ø of pre-F in enzyme-linked immunosorbent assay (ELISA). This result suggests that a low 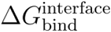 is a reliable indicator for selecting the top candidates in the subsequent design phase.

In phase 2, to further enhance antigen expression and stability, we expanded our candidate pool with additional designs featuring diverse backbones. Initial *in silico* validation using AlphaFold-Multimer ^25^ and Rosetta ^26^ confirmed that these designs were predicted to adopt the target trimeric state (Fig. 1d). From this expanded set, 200 candidates were prioritized based on two criteria: 1) a structural root mean square deviation (RMSD) < 4.0 Å between the designed and predicted models, and 2) the lowest Rosetta energy scores. The average RMSD and Rosetta energy score of the top 200 candidates were 1.31 Å and -3405.6 Rosetta Energy Unit (REU), respectively. To identify the most promising candidates for experimental validation, we performed molecular dynamics (MD) simulations followed by MM-PB(GB)SA free energy calculations to evaluate trimer stability ^27^. We also predicted the structures of top candidates using Boltz-1 ^28^ and considered its model quality metric, interface predicted TM-score (ipTM), in our selection process. Ultimately, six designs that demonstrated superior 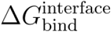 and Boltz-1 ipTM compared to p1-1 were selected for experimental validation (Fig. 1e). Seven designs, including p1-1 from phase 1, were used for experimental characterisation (Extended Data Fig. 3).

Seven constructs, including p1-1 (aRF2), were transiently expressed in Chinese hamster ovary (CHO) cells. The proteins secreted in the culture supernatant were analysed using ELISA with the monoclonal antibody, D25, to evaluate the preservation of site Ø ^29^. Immunoblotting with an anti-His antibody was also performed to confirm the expression. Among the seven constructs, three (aRF3, aRF6, and aRF7) exhibited strong binding to D25 and robust expression, whereas the others exhibited minimal or undetectable levels (Fig. 1f). We next examined protein yield after purification (Extended Data Fig. 4) and reassessed the samples using ELISA with D25 to evaluate the preservation of site Ø. Purified DS-Cav1, stabilized full length pre-F, was used as a pre-F antigen. Only aRF6 exhibited binding activity comparable to pre-F, confirming the proper presentation of the site Ø epitope after purification (Fig. 1g). Consequently, we selected aRF6 as the lead RSV vaccine antigen candidate. To further characterise the epitopes of the F protein presented on aRF6, ELISA was performed using a panel of monoclonal antibodies targeting distinct antigenic sites (Ø, I, II, III, IV, and V) (Fig. 1h, Extended Data Fig. 5). Consistent with the design, aRF6 showed exclusive binding to antibodies recognizing sites Ø, II, and V, with no or negligible binding to sites I, III, or IV, consistent with our design.

### Protective and safe immunity induced by aRF6 in preclinical models

To evaluate the immunogenicity and protective efficacy of aRF6, BALB/c mice were immunised three times with antigens formulated with aluminium hydroxide (alum) and poly(I:C), as reported previously (Fig. 2a) ^30^. Sera were collected to quantify vaccine-induced neutralising activity using a focused reduction neutralisation test (FRNT) (Fig. 2b). Sera from aRF6-immunised mice showed a measurable neutralising activity against RSV A2 (RSV A) and RSV 18537 (RSV B), although the titres were moderately lower than those elicited by the pre-F antigen. This indicates that aRF6 elicits broad nAbs with functional efficacy, despite being designed based on the RSV A2 sequence. Following the intranasal RSV A2 challenge, aRF6-immunised mice maintained their body weight, with no significant difference compared to the body weights of mice in the pre-F or uninfected groups, whereas the adjuvant-only controls began to lose weight at 6 days post-infection (dpi) (Fig. 2c). Consistently, lung viral titres were below the limit of detection in the aRF6- and pre-F-vaccinated groups, indicating sterile protection, in contrast to the high viral loads observed in the adjuvant-only controls (Fig. 2d). Histopathological analysis revealed minimal pulmonary inflammation in the aRF6 and pre-F groups, whereas the adjuvant-only group exhibited moderate-to-severe peribronchiolar and perivascular inflammatory infiltrates with marked alveolar damage, as reflected by the histological injury scores (Fig. 2e, f). These results demonstrated that aRF6 induces robust humoral immunity and confers potent protection against RSV.

**Fig. 2.**
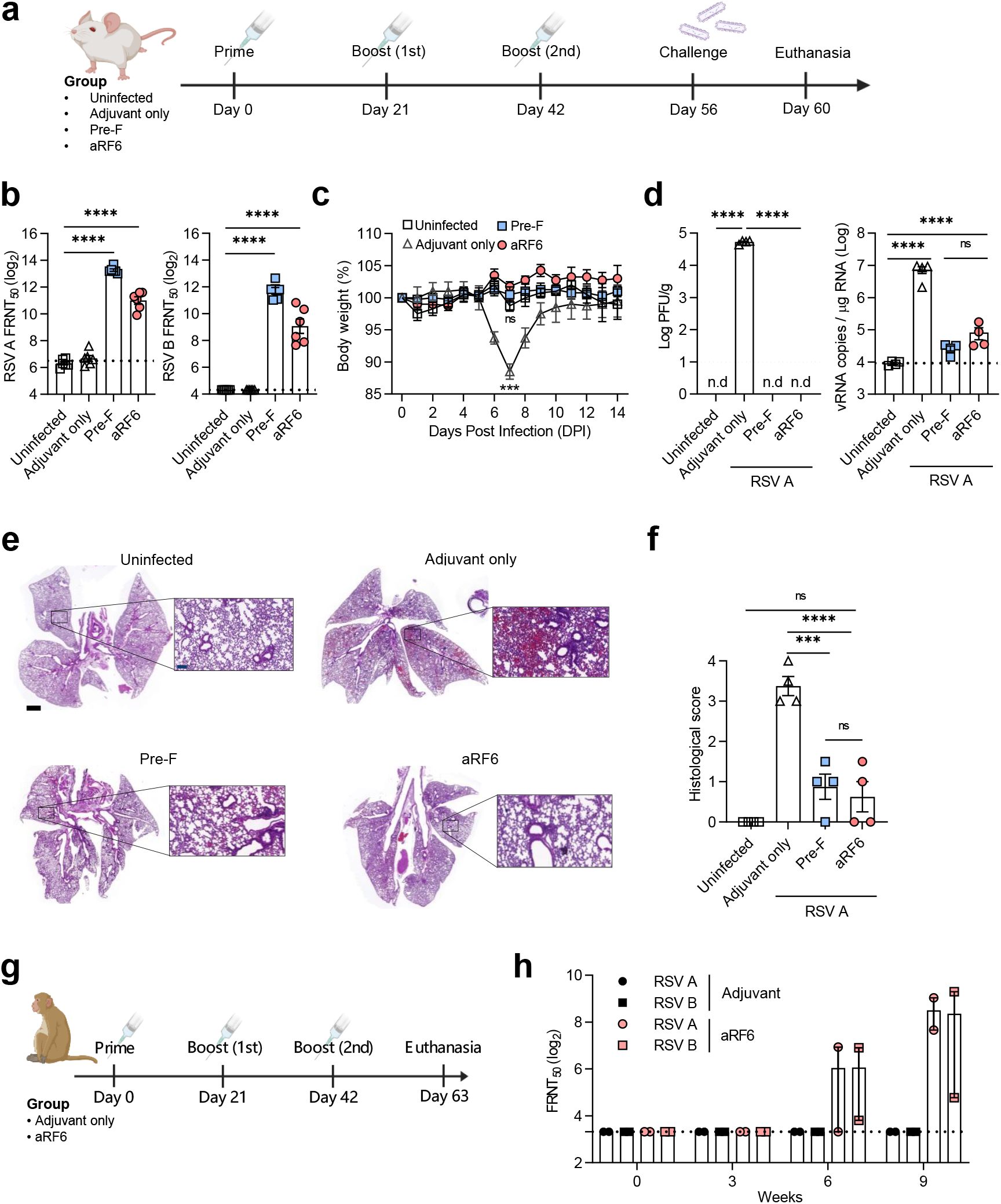
Vaccination with aRF6 elicits protective response against RSV infection *in vivo*. **a**, Schematic of the vaccination and challenge experiments. Mice were immunised three times at three-week intervals with 10 μg of the designed F protein (aRF6) formulated with 50 μg alum and 50 μg poly(I:C) adjuvants. Adjuvant-only and stabilised full-length pre-F (pre-F)-vaccinated mice served as controls. **b**, Two weeks after the final immunisation, sera were collected to measure neutralising antibody titres, expressed as the 50% focus reduction neutralisation titre (FRNT_50_), against RSV A and B prior to viral challenge (n = 6 per group). **c**, All groups were subsequently challenged with RSV A (1 × 10^6^ PFU per mouse) to assess the protective efficacy of aRF6. Body weight changes were monitored daily following viral challenge (n = 4 per group). **d**, Four days after the challenge, mice were euthanised and the lungs were harvested to quantify viral titres, including infectious virus levels and viral genomic RNA (n = 4 per group). **e**, Lung histopathology was evaluated using haematoxylin and eosin (H & E) staining, and representative images from each group are shown with low (scale bar = 1000 μm) and high magnification (scale bar = 50 μm). **f**, The pathological scores were assigned on a scale ranging from 0 to 4 (n = 4 per group). Data from BALB/c mice are representative of two independent experiments. Error bars indicate the mean ± SEM. Statistical analyses were performed using one-way ANOVA (a–e). **g**, Schematic diagram of the vaccination and challenge of non-human primates (NHPs). NHPs were randomly assigned to receive the aRF6 vaccination, with PBS acting as a control (n = 2 per group). **h**, Three doses were administered at three-week intervals, and blood samples were collected prior to each immunisation for FRNT_50_ analysis. Error bars indicate the mean ± SEM.

Building on the vaccine efficacy observed in murine models, the immunogenicity of aRF6 was assessed in NHPs (long-tailed macaque, *Macaca fascicularis*). The animals received prime immunisation, followed by two booster doses at three-week intervals (Fig. 2g). Vaccination with aRF6 induced potent RSV-neutralising titres against both RSV types A and B (Fig. 2h). Additionally, comprehensive toxicological analyses were conducted on the NHPs to evaluate the safety profile of the *de novo* antigens. Selected haematological and immunological parameters remained within established physiological ranges throughout the study (Extended Data Fig. 6)^31,32^. Furthermore, histopathological examination of five major organs— the heart, liver, kidney, spleen, and lungs —did not reveal any lesion indicative of vaccine-associated toxicity (data not shown).

### Mitigation of ERD and Th2-biased responses by aRF6 in a stringent mouse model

ERD remains a critical safety concern for the development of RSV vaccines. To assess the potential of aRF6 to induce ERD, mice were immunised with alum as the sole adjuvant, which is known to promote Th2-biased responses (Fig. 3a) ^6,33,34^. As expected, FI-RSV-vaccinated mice exhibited rapid weight loss and increased lung weight, consistent with the hallmark manifestations of ERD (Fig. 3b, c). In contrast, aRF6- and pre-F-vaccinated mice maintained stable body weights comparable to those of the uninfected controls. Histopathological examination revealed severe pulmonary pathology in the FI-RSV group, characterised by pronounced peribronchial inflammation and disruption of the alveolar septa, with extensive immune cell infiltration, leading to bronchiolar lumen obstruction. In contrast, aRF6-immunised mice exhibited minimal histopathological changes with largely preserved lung architecture, as confirmed by quantitative lung injury scoring (Fig. 3d, e). In addition, analysis of mucus production in the bronchiolar airspaces revealed significant mucus overproduction in the FI-RSV group, characterised by excessive mucus lining the apical epithelium and goblet cells of the bronchioles. Mild mucus production was observed in the pre-F-immunised group, whereas no detectable mucus accumulation was observed in the aRF6-immunised group (Fig. 3d, f).

**Fig. 3.**
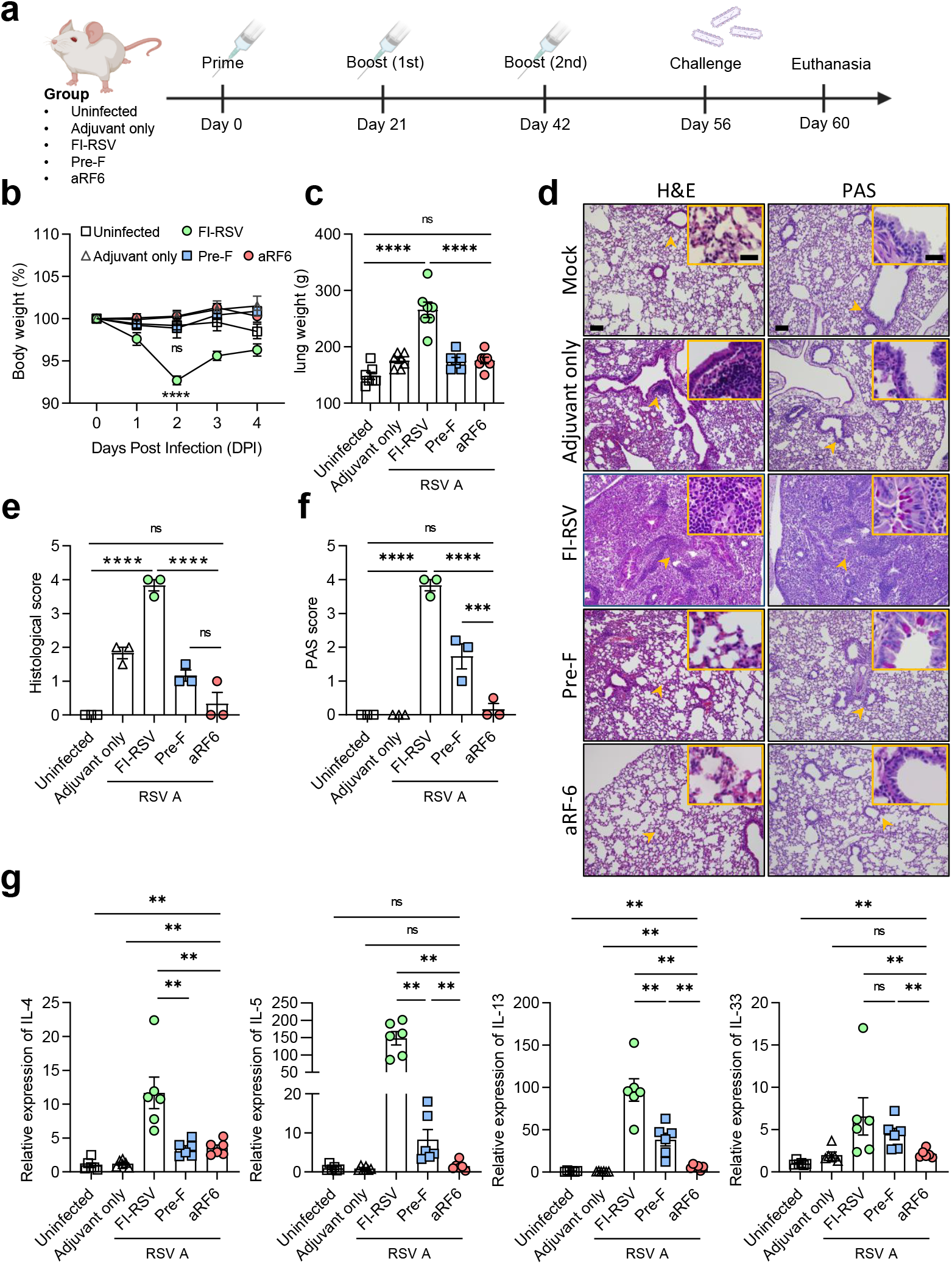
Reduced enhanced respiratory disease (ERD)–associated biomarker in a stringent mouse model. **a**, Schematic of the vaccination and RSV challenge regimens used to evaluate enhanced respiratory disease (ERD). Mice were immunised three times at three-week intervals with 10 μg of the designed F protein (aRF6) formulated with 50 μg alum. PBS-, FI-RSV-, and pre-F-vaccinated mice were used as controls. **b**, Mice were challenged intranasally with RSV A at a dose of 1 × 10^6^ PFU per mouse. Changes in body weight were monitored daily for 4 days post-infection (dpi) (n = 7 per group). **c**, At 4 dpi, the mice were euthanised and the lungs were harvested and weighed (n = 7 per group). **d**, Lung histopathology was assessed using haematoxylin and eosin (H & E) staining, and mucus accumulation was evaluated using periodic acid–Schiff (PAS) staining. Representative images from each group are shown at low magnification (scale bar, 100 μm) and high magnification (scale bar, 25 μm). **e-f**, Histopathological (e) and mucus accumulation scores (f) were assigned on a scale of 0–4 based on established criteria. (n = 3 per group). **g**, mRNA expression of Th2-type and lung injury-associated cytokines (*Il4, Il5, Il13*, and *Il33*) was quantified in the lung tissues. Cytokine expression levels were normalised to β-actin level, and fold changes were calculated relative to the uninfected group. Statistical analysis was performed using one-way analysis of variance (ANOVA) (b-f) and the Mann–Whitney U test (g). Error bars indicate the mean ± SEM.

We further analysed the expression of ERD-associated cytokines, including interleukin (IL)-4, IL-5, IL-13, and IL-33 (Fig. 3g). FI-RSV vaccination significantly elevated the mRNA levels of Th2 and lung injury-associated cytokines, which correlated with ERD. Notably, compared to the adjuvant-only and uninfected control groups, the pre-F-vaccinated group exhibited intermediate cytokine upregulation, whereas aRF6-vaccinated mice showed minimal expression, which was indistinguishable from the levels in the control groups. Collectively, these findings demonstrated that aRF6 provides robust protection without Th2 skewing or pulmonary pathology associated with ERD, offering a safety profile superior to that of pre-F in this stringent ERD model.

### Immuno-focused antibody responses and minimised scaffold immunogenicity induced by aRF6

To concentrate the immune responses on neutralising epitopes within the pre-F head domain, the aRF6 design explicitly excluded non-essential regions, thereby minimising non-nAbs (Fig. 1b). To assess the efficacy of this approach, we analysed the cross-reactivity of sera obtained from mice vaccinated with wild type (WT) F protein, pre-F, or aRF6 (Fig. 4a, left). Although all groups showed comparable levels of antibody binding to the pre-F antigen, distinct binding patterns were observed for post-F. In particular, sera from WT F-vaccinated mice exhibited the highest binding affinity to post-F, while those from the pre-F mice and aRF6-vaccinated mice exhibited intermediate binding and minimal cross-reactivity with post-F, respectively, consistent with the intended antigen design (Fig. 4a, middle). Conversely, sera from WT F- and pre-F-vaccinated mice showed negligible binding to the aRF6 antigen, whereas sera from aRF6-vaccinated mice retained robust binding to aRF6 (Fig. 4a, right). This finding indicated that antibodies induced by aRF6 vaccination were highly specific to the designed epitope display. Consequently, vaccination with aRF6 produced the highest pre-F/post-F binding ratio, significantly surpassing the ratios observed following immunisation with either the pre-F- or WT F antigen (Fig. 4b). These results suggested that aRF6 elicits a highly specific antibody repertoire that preferentially targets pre-F epitopes. To further identify the antigenic regions targeted by the vaccine-induced antibodies, we evaluated serum binding to both aRF6 and pre-F antigens. Sera from mice vaccinated with pre-F showed significantly reduced binding to aRF6 compared to homologous pre-F, indicating that a substantial proportion of antibodies were induced by the stem region of the pre-F protein (Fig. 4c). In contrast, the sera from aRF6-vaccinated mice exhibited comparable binding to both aRF6 and pre-F antigens, demonstrating that the aRF6 elicited antibodies primarily against neutralizing epitopes rather than the scaffold.

**Fig. 4.**
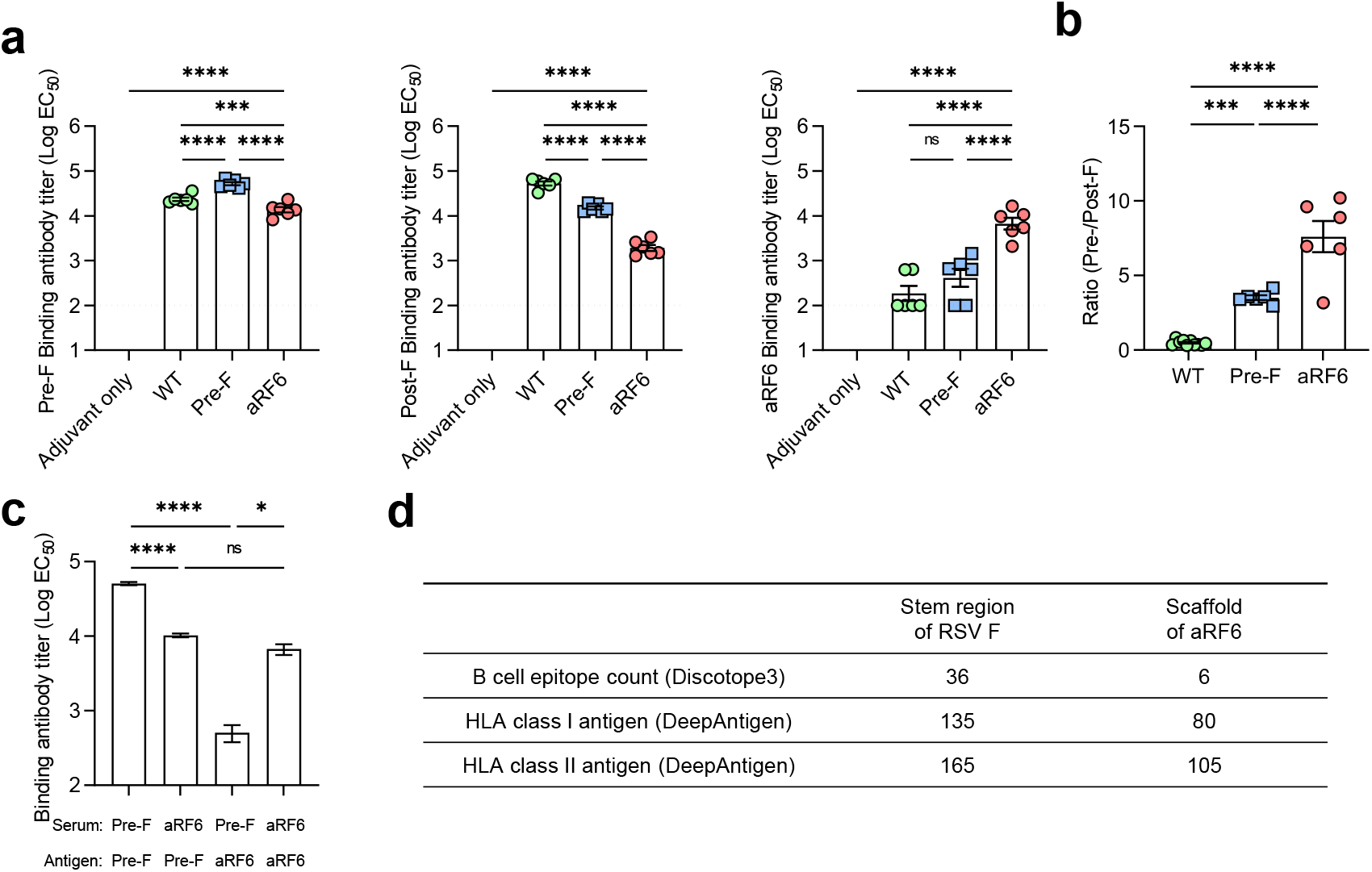
Immuno-focused profiling of serum antibody specificities after aRF6 vaccination. Serum antibody responses induced by aRF6 vaccination were characterised and compared with those elicited by the WT F protein and stabilised pre-F immunisation using an ELISA-based binding assay. **a–c**, Sera collected from mice immunised three times (n = 6 per group) were analysed for binding to recombinant pre-F, post-F, and aRF6 antigens. ELISA plates were coated with 1 μg/mL of each antigen and incubated with serially diluted serum samples, followed by detection with horseradish peroxidase (HRP)-conjugated anti-mouse IgG. Binding curves were generated by plotting absorbance values against serum dilution, and EC_50_ values were calculated using nonlinear regression with a four-parameter logistic model in GraphPad Prism. a, Binding to pre-F (left), post-F (middle), and aRF6 (right) antigens. b, Ratio of pre-F to post-F binding calculated for each immunisation group. c, Antigen-dependent cross-binding of serum antibodies to pre-F and aRF6 antigens. Statistical analyses were performed using one-way analysis of variance (ANOVA). Error bars indicate the mean ± SEM. d, The B cell and MHC loading probability of the stem region of the RSV F protein and scaffold region of aRF6 were predicted using Discotope-3.0 and deepAntigen, respectively.

Computational assessments of immunogenicity were performed to validate the experimental results. Discotope-3.0 revealed that the *de novo* scaffold of aRF6 had significantly lower B cell epitope potential than the native RSV F stem, indicating the successful replacement of the stem region to limit non-neutralising antibody induction (Fig. 4d). Additionally, DeepAntigen analysis indicated a lower major histocompatibility complex (MHC)-loading probability for the aRF6 scaffold than for the native structure. These *in silico* predictions support the idea that the aRF6 design effectively dampens scaffold-specific immunity, thereby focusing on the immune response toward the target pre-F epitopes. Collectively, these data indicated that the antibody response induced by aRF6 is predominantly focused on conserved neutralising epitopes (Ø, II and V), rather than the *de novo* scaffold, thereby confirming the minimised immunogenicity of the scaffold region.

### Cryo-electron microscopy (EM) analysis reveals an aRF6 structure closely matching the designed model

We previously engineered aRF6 to assemble as a trimer and observed that the D25 antibody, which recognizes the trimeric head of pre-F, also binds aRF6, suggesting that aRF6 adopts a trimeric structure similar to that of pre-F. To verify the trimeric nature of aRF6 and assess its structural correspondence with the pre-F head, we determined the aRF6 structure using cryo-EM.

Two-dimensional class averages of the extracted particles revealed a tetrahedral morphology that closely matched the 2D templates generated by the trimeric design model (Extended Data Fig. 7).

The resulting cryo-EM reconstruction also displayed a tetrahedral assembly with clear C3 symmetry, and the designed model fit the map with excellent agreement, prompting further refinement, considering the well-resolved α-helical features (Fig. 5a). The loop region at the top of the head was not resolved, possibly because of conformational flexibility, and a part of the C-terminus was missing, consistent with the presence of a disordered C-terminal His tag used for purification.

**Fig. 5.**
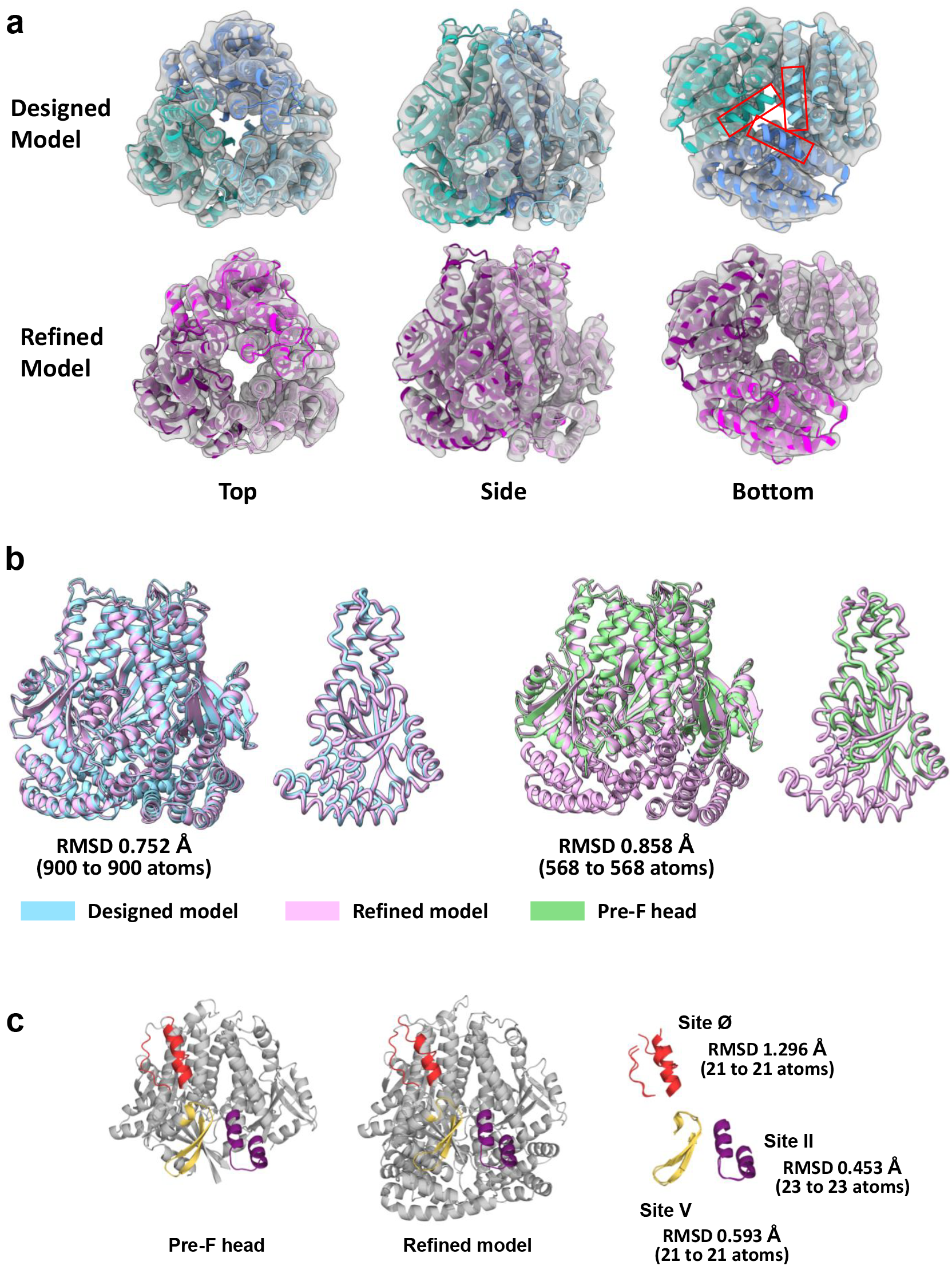
Comparative structural analysis of the aRF6 antigen determined using cryo-EM. **a**, Cryo-EM map of the aRF6 antigen with the fitted designed model and refined model. **b**, Structural comparison of the designed model and refined model, as well as the pre-fusion (pre-F) head domain and the refined model. The designed model is shown in light blue, the refined model in light purple, and the pre-F head domain in light green. **c**, Comparison of the preserved sites Ø, II, and V between the pre-F head and the refined model. Site Ø is shown in red, site II in purple, and site V in yellow.

We performed structural superimposition to evaluate the structural similarity between the models. The refined aRF6 model showed high structural similarity to the designed model, with an RMSD of 0.752 Å, and also aligned closely with the pre-fusion head-domain template used in the initial design, with an RMSD of 0.858 Å (Fig. 5b). Crucially, sites Ø, II, and V were structurally preserved as designed, with RMSD values of 1.296 Å, 0.453 Å, and 0.593 Å, respectively (Fig. 5c). The higher RMSD for site Ø possibly reflects flexibility in the loop region at the top of the head, whereas site II and V showed minimal deviation. These results demonstrated that aRF6 adopts a trimeric architecture that closely recapitulates the pre-F head structure, indicating that the key neutralising epitopes are well preserved.

## Discussion

RSV vaccine development has progressed substantially through structure-based approaches that exploit potent neutralising epitopes presented by pre-F ^10^. Pre-F elicits substantially higher neutralising antibody titres than post-F, and this discovery has enabled the development of currently licensed vaccines ^13,14,35,36^. However, critical gaps remain: vaccines specifically licensed for high-risk young infants remain unavailable, and ERD continues to be a cause of concern in defined settings. Recent clinical safety signals associated with the increased RSV-associated hospitalizations in infants receiving pre-F mRNA vaccine candidates suggest that stabilized full-length pre-F designs may not represent an immunologically optimal strategy for this age group. To address these limitations, we developed a *de novo* antigen using an AI-based protein design that incorporated an immuno-focusing strategy. Rather than stabilizing entire native pre-F protein, which retain non-essential domains, including the stem region and a heterologous trimerization domain that can promote non-neutralising antibody responses and excessive Th2-skewed immunity, we preserved only the three critical neutralising epitopes (sites Ø, II, and V) on an AI-designed scaffold engineered for minimal immunogenicity. This precision design strategy helps transcend the problems associated with conventional protein stabilization by selectively exposing immunologically favourable epitopes, while suppressing off-target immunogenicity.

Of eleven antigen candidates, three successfully presented site Ø, with aRF6 demonstrating exceptional fidelity to design intent by preserving all three neutralising epitopes (sites Ø, II, and V). The successful derivation of a lead candidate by minimal screening highlights the efficiency of combining modern deep learning-based protein design methods with physics-based MD simulations. Our results demonstrated that MD simulations efficiently evaluate the trimeric stability of the candidates by verifying the integrity of inter-protomer interfaces and quaternary structure under physiological conditions. This integrated pipeline illustrates how AI-driven protein design is complemented by physics-based validation and how it substantially reduces empirical trial and error, enabling the rapid identification of antigens with high structural fidelity. In murine immunisation studies, aRF6 elicited robust neutralising antibody responses that provided complete protection against RSV A and RSV B challenges, although they were modestly lower than those provided by pre-F. The reduced neutralising antibody titres in aRF6-immunised mice may reflect the absence of TLR4-stimulating motifs, including site IV, present on full-length pre-F but absent in aRF6 ^37,38^. Critically, nonclinical assessment of NHP revealed not only the formation of antigen-specific neutralising antibodies but also the absence of pathological alterations or systemic toxicity following antigen administration.

In stringent murine models designed to maximize ERD development, aRF6 vaccination resulted in minimal pulmonary pathology, which was substantially lower than that observed with pre-F- or FI-RSV, and it approached the levels observed in uninfected or alum-only controls. Th2-type immune responses and lung injury were markedly attenuated in the aRF6-vaccinated mice, indicating a pronounced reduction in ERD-associated immunopathology. This reflects two key features of the design. First, aRF6 elicits highly selective pre-F-directed antibody responses with markedly reduced binding to post-F and non-neutralising epitopes, thereby focusing on humoral immunity triggered by protective epitopes while suppressing off-target antibody formation. Second, the immuno-focused antigen design minimises cell-mediated immune reactivity because aRF6 exposes only three essential epitopes rather than the full-length pre-F protein, thereby reducing the breadth of T cell recognition. This combined achievement of robust protection and enhanced safety supports AI-based *de novo* antigen design as a powerful approach to mitigate the excessive Th2-skewed immune responses observed with conventional pre-F immunogens.

These findings establish that AI-based *de novo* antigen design employing an immuno-focusing approach can elicit potent neutralising activity against RSV infection, while substantially mitigating ERD-associated risks, which is a critical requirement for vaccines intended for infants. Our aRF6 immunogen exemplifies how the selective display of functional epitopes, except non-neutralising epitopes, coupled with the suppression of off-target immune responses to AI-based scaffolds, resolves the safety limitations that have constrained the clinical translation of full-length pre-F designs. Beyond RSV, this approach has the potential to mitigate the conserved mechanisms of ERD documented for various respiratory viruses, such as influenza, SARS-CoV-2, and measles ^39-42^. This immuno-focusing strategy provides a generalizable design framework that may improve vaccine safety across the respiratory viral landscape. Integration with diverse vaccine platforms, including mRNA, recombinant viral vectors, and nanoparticle formulations, may further enhance immunogenicity while preserving the safety advantages demonstrated herein, positioning these immunogens as foundational candidates for clinical development in high-risk populations with potential applicability beyond the RSV context.

## Methods

### Initial antigen design phase

RF diffusion ^23^ was used to generate novel protein backbone structures while preserving the sites Ø and V of the pre-fusion structure (PDB: 4JHW ^10^ of RSF F protein (residue numbers 51-97 and 149-305). Designs were generated with C3 symmetry to match the inherent C3 symmetry of the RSV F protein trimer. Oligomer contact guiding potentials were used during the generation processes to strengthen the interprotomer interactions and generate ten backbone designs. For each backbone, 100,000 sequences were generated using ProteinMPNN ^24^ with a sampling temperature of 0.3 and the v_48_020 parameter set. The top 100 sequences with the lowest ProteinMPNN scores, the negative log likelihood of a generated sequence for a given structure, were selected from the 100,000 generated sequences. In total, 1,000 candidate sequences were advanced for stability and free-energy calculation analyses.

The trimer structures of the designed sequences were cross-validated using AlphaFold-multimer ^25^. For each candidate sequence, AlphaFold-multimer predicted 25 model structures. Among the model structures, the most confident model structure with the highest plDDT was chosen as the representative structure of the designed sequence. The RMSD between the designed backbones and their AlphaFold-multimer-predicted counterparts was calculated using US-align ^25^. The designs with RMSD > 4.0 Å were filtered out and the stabilities of the remaining complexes were assessed using the Rosetta energy function ^26^. The top 100 designs with the lowest Rosetta energies were selected for further analysis. For these 100 designs, three independent 100 ns molecular dynamics (MD) simulations were performed using the most confident predicted structure from AlphaFold-multimer with the ff14SB force field ^43^ and TIP3P water model ^44^. The interfacial binding free energy of a protomer to the remaining dimer was calculated using the Molecular Mechanics/Poisson-Boltzmann Surface area (MM/PBSA) and Molecular Mechanics/Generalized Born Surface Area (MM/GBSA) methods using the final 10 ns of each trajectory ^27^. For convenience, designs whose N- and C-termini were not exposed were excluded. The four most stable designs with the lowest interfacial binding affinities were selected for experimental validation.

### Expanded design phase

To explore a wider backbone configurational space, additional 90 backbones were generated and 100 sequences were selected per backbone. Backbones with buried N-terminal- and C-terminal regions were excluded from the analysis. The top 200 designs were selected using the identical RMSD and Rosetta energy criteria as the initial design phases. For the selected 200 designs, MD simulations were performed and their interfacial binding energies were calculated. Designs with interfacial binding energies lower than the initial phase hits were maintained. For further *in silico* cross-validation, the trimer structures of the designs were predicted using Boltz-1 ^28^, and the designs with ipTM < 0.7 were filtered out. Among the remaining candidates, six most stable designs with the lowest interfacial binding free energies were selected for experimental validation.

### Prediction of B cell antigenicity

Structure-based B cell epitope prediction was performed for aRF6 and RSV F proteins using DiscoTope-3.0 ^45^. Chain A of the AlphaFold-Multimer predicted that the trimeric structures would act as inputs. A strict prediction score threshold of 1.5 was applied to identify B cell epitopes, and the total number of predicted epitope residues was quantified for each protein.

MHC loading probabilities were predicted using deepAntigen ^46^. Although the model supports structural integration, the predictions in this study were made using only sequence inputs. All overlapping 8–11-mer peptides derived from aRF6 and RSV F proteins were evaluated against a binding probability threshold of 0.5. The number of identified binders was counted across a reference set of 27 MHC class I allele combinations (HLA-A and HLA-B) and 27 MHC class II allele combinations (HLA-DR, HLA-DQ, and HLA-DP).

### Cell and virus propagation

HEp-2 cells (KCLB, 10023, Korean Cell Line Bank, Seoul, Korea) were maintained in minimum essential medium (MEM; Gibco, 11095-080) supplemented with 100 U/mL penicillin-streptomycin (Gibco, 15140-122) and 10% (v/v) foetal bovine serum (FBS; Gibco, 16000-044). The CHO-S (Gibco, A29133) and Epxi293 (Gibco, Waltham, MA, USA, A41249) cells were purchased from Gibco. RSV strains A2 (ATCC, VR-1540) and B 18537 (ATCC, VR-1580) were propagated in HEp-2 cells. Viral titres were determined using plaque assays.

### Mice

Six-week-old female BALB/c mice were purchased from Orient Bio (Seongnam City, South Korea) and housed in a biosafety level 2 (BSL-2) facility at the Korea Research Institute of Chemical Technology (KRICT), Daejeon, South Korea. All protocols were approved by the Institutional Animal Care and Use Committee (Protocol ID 8A-M10, IACUC ID 2025-8A-05-01, and 2025-8A-08-03). For immunisation, mice were intramuscularly injected with 10 µg constructs or control antigens adjuvanted with 50 µg of 2% Alhydrogel (Alum, Vac-alu, InvivoGen, San Diego, CA, USA) and 50 µg poly I:C (Sigma-Aldrich, St. Louis,MO, USA, P0913) three times with an interval of three weeks between each injection. To challenge the vaccinated mice, isoflurane was used for sedation and the mice were intranasally inoculated with 1 × 10^6^ plaque-forming units (PFU) of RSV A2.

### Protein expression and purification

DS-Cav1 (PDB: 6VKD) was used as the reference antigen for the stabilized pre-fusion RSV F protein. The gene sequences of the control antigen and designed constructs were synthesised by Integrated DNA Technologies (IDT) and cloned into pcDNA3.1^(+)^ (Invitrogen, Carlsbad, California, USA) with a 6× His-tag at the C-terminus. All recombinant proteins were expressed in CHO-S or Expi293 cells according to the manufacturer’s instructions. In brief, cells in suspension culture were transfected with clones encoding each recombinant protein and incubated at 37°C in an atmosphere of 8% CO_2_ with agitation at 120 rpm on a shaker (20 mm shaking diameter). The culture supernatant was harvested via centrifugation at 2,000 × *g* for 20 min before the cells reached 80% confluence. Then, the collected supernatant was filtered through a 0.8-µm filter and loaded onto Ni Sepharose 6 FF (Cytiva, MA, USA) pre-equilibrated with buffer A (500 mM sodium chloride and 20 mM HEPES pH 7.5). The bound proteins were eluted using a linear gradient buffer B (500 mM sodium chloride, 20 mM HEPES pH 7.5, and 500 mM imidazole). The proteins were further purified via size-exclusion chromatography using a HiLoad 26/600 Superdex 200 pg column (Cytiva) equilibrated with buffer A. The fractions eluted from the major peaks were collected and concentrated to 0.6 mg/mL.

### Immunoblotting

The media containing recombinant proteins were separated via electrophoresis on denaturing 10% sodium dodecyl sulphate-polyacrylamide gel and transferred to polyvinylidene difluoride membranes (1620177, Bio-Rad, Hercules, CA, USA). Membranes were blocked with 5% skim milk (LPS solution, skI500) in Tris-buffered saline containing 0.1% Tween 20 (TBS-T) for 1 h at 24°C. Subsequently, the target proteins were detected using an anti-His antibody conjugated with HRP (Bethyl Laboratories, Montgomery, Texas, USA, a190-114p) and visualised using chemiluminescence reagents (34580, Thermo Fisher Scientific).

### ELISA

Whether the expressed recombinant proteins contained epitopes consistent with the intended design was verified. Maxibinding flat-bottom 96-well immunology plates (32296, SPL Life Sciences, Pocheon City, South Korea) were coated with 0.2 μg/mL D25 monoclonal antibody (01-07-0120, Cambridge Biologics, Cambridge, MA, USA) in phosphate buffered saline (PBS) and incubated overnight at 4°C. The plates were blocked with 2% skim milk in TBS-T for 2 h at 24°C. Serially diluted expression media containing recombinant proteins was added to the plates and incubated for 2 h at 24°C on a rocker. The captured recombinant proteins were detected using an anti-His antibody conjugated with HRP.

To evaluate the binding affinity to site Ø of F protein-specific mAb, purified recombinant proteins and WT F (11049-V08B, Sinobiological Inc, Beijing, China) and post-F (RSF-V52H6, Acro biosystem, Newark, DE, USA) were serially diluted two-fold from the initial concentration of 1 µg/mL and used to coat immunology plates overnight at 4°C. After blocking with 2% (w/v) skim milk in TBS-T, the site Ø of each recombinant protein was detected using D25 mAb (1:2,000) and HRP-conjugated anti-human IgG.

The plates were washed thrice with TBS-T between each step, and colour development was initiated by adding 1-Step Ultra TMB-ELISA substrate solution (34028, Thermo Fisher Scientific). The reaction was stopped using 0.5 M HCl. Absorbance was measured at 450 nm.

To determine the binding Ab titre induced by vaccination, immunology plates were coated with 1 μg/mL of pre-F, post-F, or aRF6 protein (DAG-WT1180, Creative diagnostics). After blocking with 2% skim milk in TBS-T, three-fold dilutions (1:100) of each protein was added to plates and incubated for 1 h at 24°C on a rocker, followed by the addition of HRP-linked anti-mouse IgG (1:5,000; Cell Signaling Technology, Danvers, MA, USA). The colour development and measurement procedures were the same as those described above. Binding curves were generated by plotting absorbance values against serum dilution, and EC_50_ values (half-maximal effective concentration) were calculated using nonlinear regression with a four-parameter logistic model in GraphPad Prism.

### Neutralising Ab titration

The neutralising ability of the collected serum samples was determined using a FRNT. Briefly, HEp-2 cells were seeded in a 96-well plate one day before the assay. Heat-inactivated mouse sera were serially diluted in MEM supplemented with 2% FBS and incubated with 500 focus forming units (FFUs) of RSV A2 or RSV 18537 for 90 min at 37°C in a final volume of 100 μL. This mixture was transferred to plates containing HEp-2 cells and incubated for 30 h at 37°C in an atmosphere of 5% CO_2._ The cells were washed with cold PBS, fixed, and then stained with an RSV N-specific antibody (GeneTex, GTX636648) and secondary HRP-conjugated goat anti-rabbit IgG (GeneTex). Foci signals were developed using an insoluble TMB substrate (Promega, Madison, WI, USA) and counted using an ImmunoSpot reader (CTL, Shaker Height, OH, USA).

### Reverse transcription-quantitative polymerase chain reaction (RT-qPCR)

Viral RNA and gene expression were quantified using RT-qPCR on a QuantStudio 3 real-time PCR system (Applied Biosystems, Foster City, CA, USA). The expression of the RSV F gene and *Il4* was detected using the one-step RT-qPCR with PrimeScript III RT-qPCR mix (Takara Bio, Shiga, Japan) and TaqMan probe-based assays. For RSV F, a custom probe-based assay (Integrated DNA Technologies, Coralville, IA, USA) was used with the following oligonucleotides: forward primer 5′-AACAGATGTAAGCAGCTCCGTTATC-3′, reverse primer 5′-GATTTTTATTGGATGCTGTACATTT-3′, and fluorescently labelled probe 5′-FAM-TGCCATAGCATGACACAATGGCTCCT-TAMRA-3′. *Il4* expression was quantified using a pre-designed TaqMan assay (Integrated DNA Technologies, Coralville, IA, USA).

For *Il5, Il13, Il33*, and *Gapdh*, RT-qPCR was performed using TB Green Premix Ex Taq (Takara Bio) with CYBR Green detection and the following primer sets: *Il5* (forward: 5′-GATGAGGCTTCCTGTCCCTACT-3′, reverse: 5′-TGACAGGTTTTGGAATAGCATTTCC-3′); *Il13* (forward: 5′-AACGGCAGCATGGTATGGAGTG-3′, reverse: 5′-TGGGTCCTGTAGATGGCATTGC-3′); *Il33* (forward: 5′-CTACTGCATGAGACTCCGTTCTG-3′, reverse: 5′-AGAATCCCGTGGATAGGCAGAG-3′); and *Gapdh* (forward: 5′-TCTCCTGCGACTTCAACA-3′, reverse: 5′-TGTAGCCGTATTCATTGTCA-3′).

### Lung histopathology

Five days after the virus challenge, the mice were euthanised and subjected to cardiac perfusion with cold PBS. Whole lung tissues were harvested and fixed in 10% neutral-buffered formalin for 48 h at room temperature, followed by paraffin embedding and microtome sectioning at 4-μm thickness. Lung tissue damage was evaluated using H & E staining, whereas mucus production was assessed using PAS staining. Whole-slide images were acquired using a Pannoramic MIDI slide scanner (3DHISTECH, Budapest, Hungary), and high-resolution microscopic images were obtained using an Olympus light microscope. Histopathological evaluation focused on the perivascular and peribronchial regions to assess inflammatory immune cell infiltration, as well as on the alveolar regions to evaluate pulmonary tissue architecture and structural integrity. Histological changes were quantified using a four-point severity scoring system. Inflammatory cell aggregation and interstitial pneumonia were scored as follows: 1 = normal naïve lung architecture; 2 = slight and occasional inflammatory cell aggregation; 3 = moderate inflammatory cell infiltration; and 4 = moderate to severe inflammatory pathology. For analysing mucus accumulation, PAS-stained sections were evaluated and scored based on mucus production severity as follows: 1, no detectable mucus; 2, rare presence of mucus; 3, moderate mucus accumulation; and 4, severe mucus production within the airways.

### Immune response to vaccine and toxicity evaluation in NHP

To assess the efficacy and potential toxicity of the vaccine formulation, a study was performed on long-tailed macaques (*Macaca fascicularis*). All NHP experiments were performed at the Korea Institute of Toxicology (KIT) and the IACUC protocol was followed (IAC-25-01-0048-0068). The vaccine formulation consisted of 100 g of antigen combined with 250 g of aluminium hydroxide (alum) adjuvant per dose. Four animals were randomly assigned to one of two groups: a vaccinated group (n = 2) or a negative control group (n = 2). Animals in the vaccinated group received injections of the vaccine formulation into the quadriceps muscles at weeks 0, 3, and 6. The negative control group was administered an equivalent volume of HEPES buffer, according to the same schedule. All animals were immunised three times at three-week intervals. Throughout the study, toxicity was comprehensively evaluated on the basis of daily clinical observations and weekly body weight measurements. Clinical pathology assessments were conducted every three weeks until the end of the study. Additionally, serum samples were collected before each vaccination to assess neutralising activity using the FRNT assay.

### Cryo-EM grid preparation and data acquisition

Purified aRF6 was applied to a freshly glow-discharged holey carbon-copper grid (Quantifoil R 1.2/1.3 300-mesh; EMS). Following the 3.0-s blotting step, the blotted grids were vitrified in ethane cooled by liquid nitrogen with a Vitrobot Mark IV (Thermo Fisher Scientific). Vitrification was performed at 4°C and 100% humidity. Screening for aRF6 was performed using Glacios (Thermo Fisher Scientific) operating at 200 kV. Automated cryo-EM data acquisition was performed on Glacios (Thermo Fisher Scientific) at an acceleration voltage of 200 kV using the EPU software (Thermo Fisher Scientific). The images were captured using a Falcon IV direct electron detector (Thermo Fisher Scientific). In total, 2,045 micrographs were acquired at a defocus range of -1.4 to - 2.4 µm, at a nominal magnification of 120,000×, corresponding to a physical pixel size of 0.85 Å. Each image was dose-fractionated into 40 frames with a total exposure time of 9.31 s, resulting in a total dose of 48.28 electrons per Å^2^.

### Cryo-EM data processing and model refinement

All data processing was performed using the CryoSPARC software (version 4.7.1)^47-49^. Automatic particle picking was performed using a Template Picker, yielding 4,481,268 particle picks from 2,045 micrographs. After inspecting and manually curating the picks, 745,067 particles were selected and extracted. These particles were used as the initial particle set and subjected to multiple rounds of 2D classification, resulting in 5,017 high-quality particles. An *ab-initio* reconstruction was performed using these particles. The initial map was refined using 3D refinement with C3 symmetry, followed by non-uniform refinement to obtain the final map.

Atomic models of aRF6 were constructed by fitting the predicted structure to a cryo-EM density map using UCSF ChimeraX (version 1.10) ^50^. The fitted coordinates were subjected to real-space refinement using PHENIX (version 1.21.2.5419) ^51^. Manual inspection and local adjustments of the model were performed using Coot (version 0.9.8.95) ^52^ to improve the geometry and model–map agreement. The final refined model exhibited good agreement with the cryo-EM map with a model-to-map correlation coefficient (CC_mask) of 0.7787, as assessed using PHENIX.

### Statistical analysis

All experiments were performed at least three times. All data were analysed using the GraphPad Prism 8.0 software (GraphPad Software, San Diego, CA, USA). Statistical significance was set at *P* < 0.05. The specific analytical methods are described in the figure legends.

## Acknowledgments

This research was conducted under Project No. KK2633-20 (A Study on the Convergence Platform for Infectious Disease Diagnosis and Prevention) supported by the Korea Research Institute of Chemical Technology (KRICT) and by a National Research Foundation of Korea (NRF) grant funded by the Ministry of Education, Science, and Technology (MIST) of the Korean government (RS-2023-002085678 and 2021M3A9G8025599). This study was supported by the National Research Council of Science and Technology (CRC22024-700) and KRIBB Research Initiative Program (KGM9952623 and KGM1062622). The use of the cryo-EM facilities of the NEXUS consortium was supported by a National Research Foundation of Korea grant RS-2024-00440289. We thank Core Research Facility & Analysis Center at KRIBB for assistance with cryo-EM and Global Science experimental Data hub Center (GSDC) at Korea Institute of Science and Technology Information (KISTI) for computing resources and technical support.

## Author contributions

Conceptualization: W.Y.H., J.S., H.-C.S., J.K., and Y.-C.K. ; Methodology: W.Y.H., J.S., J.C., K.B.K., H.-S.K., H.N.S., J.L., H.-C.S., J.K., and Y.-C.K. ; Investigation: W.Y.H., J.S., J.C., K.B.K., H.-S.K., G.Y.Y., D.Y.K., M.-R.C., J.S.L., S.P., S.K.L., S.j.K., B.K., and H.-C.S. ; Resources: H.N.S., J.L., H.-C.S., J.K., and Y.-C.K., ; Data curation and analysis: W.Y.H., J.S., J.C., K.B.K., H.-S.K. H.N.S., J.L., H.-C.S., J.K., and Y.-C.K., ; Writing–original draft: W.Y.H., J.S., J.C., and K.B.K. ; Writing–review and editing: D.-G.A., K.-D.K., C.K., H.N.S., J.L., H.-C.S., J.K., and Y.-C.K. ; Visualization: W.Y.H., J.S., J.C., K.B.K, H.-C.S., J.K., and Y.-C.K. ; Supervision : H.N.S., J.L., H.-C.S., J.K., and Y.-C.K. ; Funding acquisition H.-C.S., J.K., and Y.-C.K.

## Data availability

The cryo-EM maps and atomic coordinates have been deposited under the following accession codes; EMDB (EMD-68780) and PDB (22YS), respectively. Supplementary Information is available for this paper.

## Extended figure legends

**Extended Fig. 1. The stability and energy metrics of designs for the initial design phase**. Left: Scatter plot of RMSD values between the designed and predicted structures and Rosetta scores of the predicted structures. The selected 100 designs are coloured blue. Right: Scatter plot of interfacial binding affinity calculated using MMGBSA and MMPBSA. For convenience, we excluded designs, N- and C-termini of which were not exposed. The candidates selected for experimental validation are in red.

**Extended Fig. 2. Initial screening of *de novo*-designed proteins *in vitro***. Initially, five proteins in phase 1 were expressed and recovered from the medium. Their expression levels were quantified by immunoblotting with an anti-His antibody. The fidelity of the design intent was confirmed using ELISA with the D25 antibody.

**Extended Fig. 3. Sequence alignment of *de novo*-designed RSV antigen and native RSV F protein**.

**Extended Fig. 4. Purification of the *de novo* constructs**. *De novo* constructs were purified using Ni–NTA affinity chromatography, and the purified constructs were subsequently detected by immunoblotting using an anti-His antibody. This experiment was repeated twice.

**Extended Fig. 5. Antigenic profiling of aRF6. a**, The binding activity of aRF6 was assessed using ELISA with a panel of site-specific monoclonal antibodies targeting RSV F antigenic sites Ø (nirsevimab), I (131-2a), II (palivizumab, motavizumab), III (MPE8), IV (clesrovimab), and V (suptavumab). **b**, Binding activity was quantified as the area under the curve (AUC). The experiments were repeated twice, and representative data are shown and presented as mean ± SE.

**Extended Fig. 6. Clinical chemistry and haematological safety assessment of aRF6 vaccination in non-human primates**. Clinical chemistry and haematological parameters were evaluated in non-human primates (NHPs) vaccinated with aRF6 or PBS control, as described in Fig. 2g–h. Serum samples were collected one day prior to immunisation and at the indicated time points following vaccination. Results are shown as **a**, biochemical parameters; **b**, leukocyte- and thrombocyte-related haematological parameters; and **c**, erythrocyte-related haematological parameters. Each group consisted of two macaques (n = 2 per group). Black circles indicate PBS-treated controls and filled red squares indicate aRF6-vaccinated NHPs. Gray shaded areas represent the established physiological reference ranges for healthy macaques. Error bars indicate the mean ± SEM.

**Extended Fig. 7. 2D class average from selected particles and created 2D template from designed aRF6 model. a**, In total, 5,017 particles were classified into 21 2D classes using cryoSPARC. **b**) The aRF6 model was converted into a density map using ChimeraX and used to generate 2D templates in cryoSPARC.

