## Extended data for "*De novo* design of a safe and potent respiratory syncytial virus immuno-focusing antigen"

**Extended Data Fig. 1** The stability and energy metrics of candidates for initial design phase.

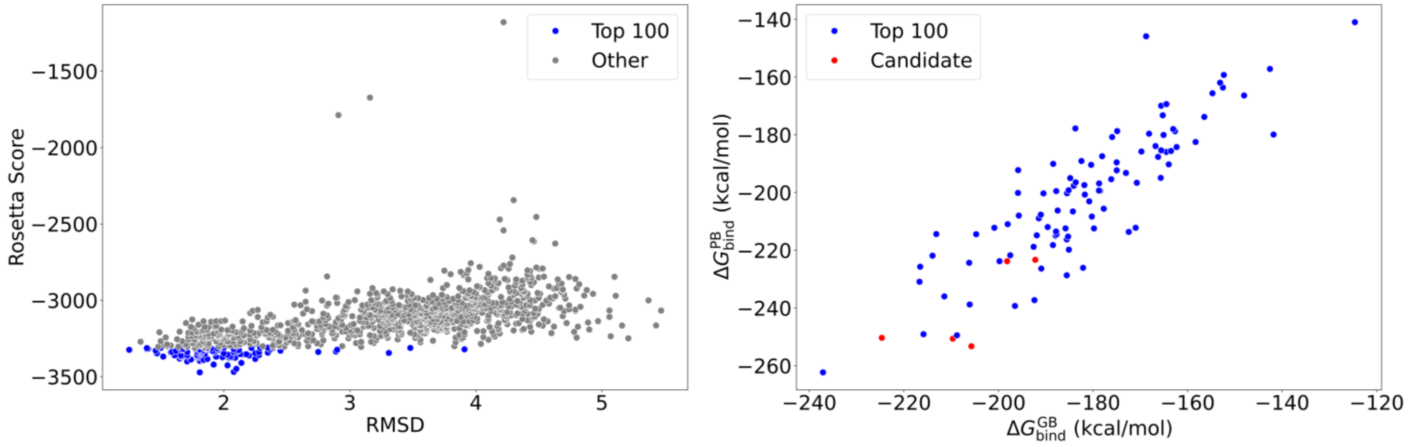

### Extended Data Fig. 2

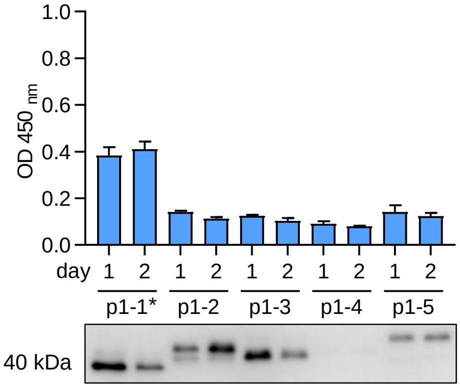


*p1-1 (aRF2)

### Extended Data Fig. 3 Sequence alignment of de novo designed RSV antigen and native RSV F protein for phase 1 and 2

**Phase 1**

Sequences will be opened after publication

**Phase 2**

Sequences will be opened after publication

### Extended Data Fig. 4

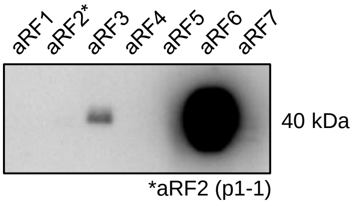


### Extended Data Fig. 5

**a**

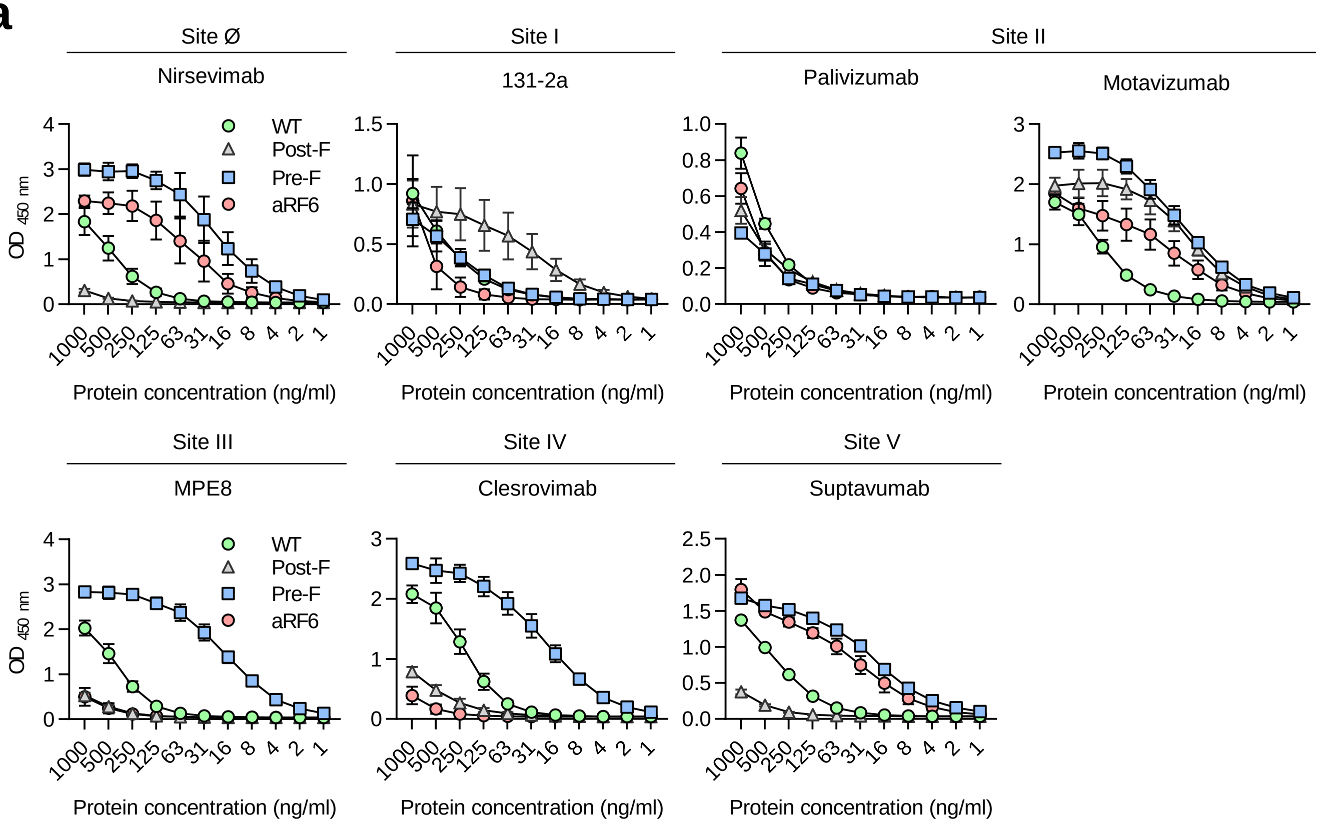


**b**

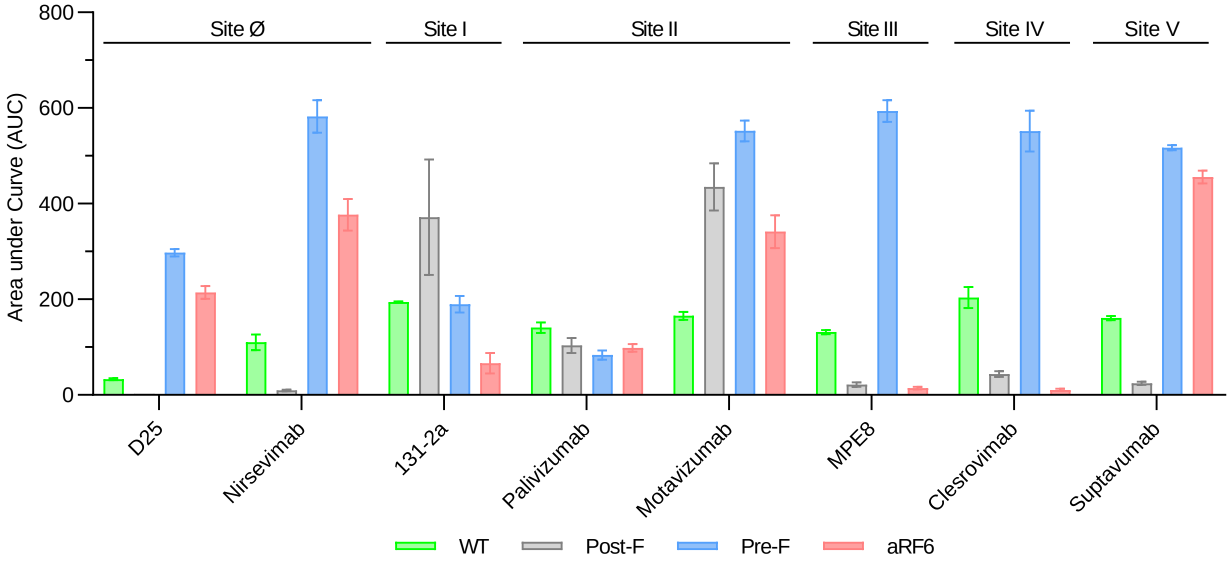

#### Extended Data Fig. 6

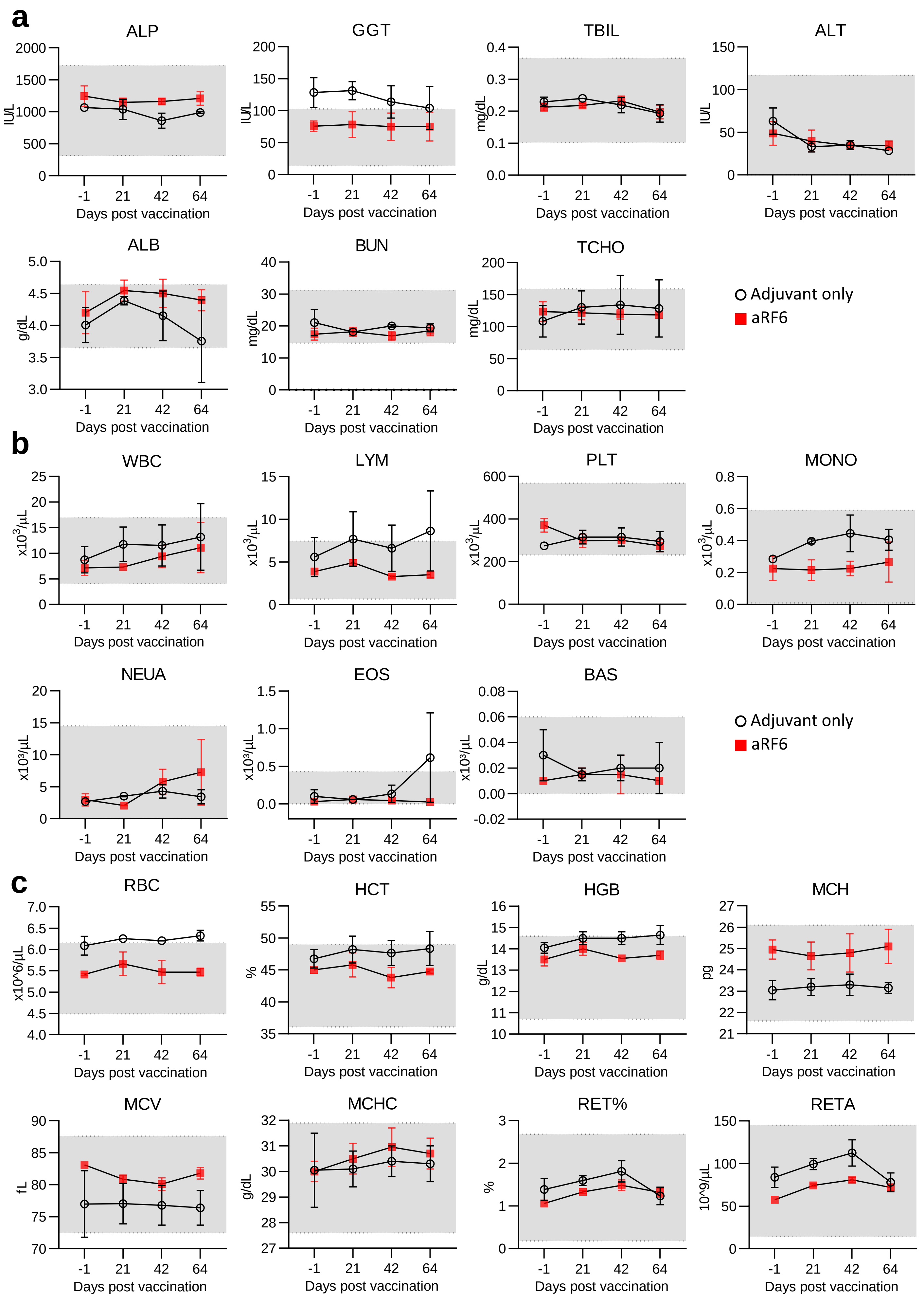

#### Extended Data Fig. 7

**a**

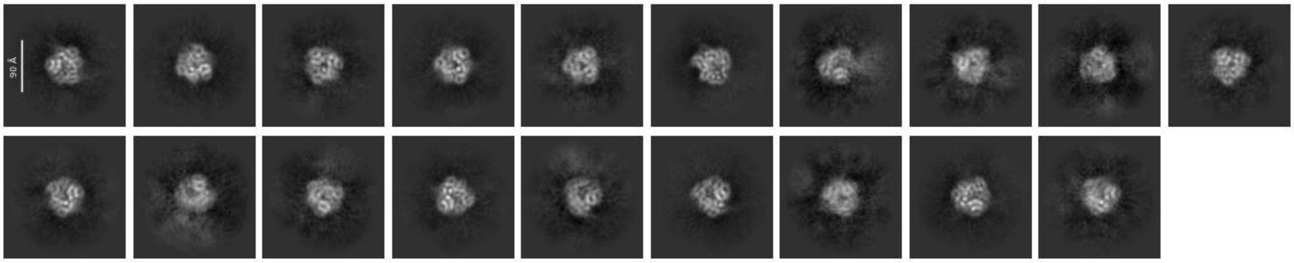


2D class average from the extracted particles

**b**

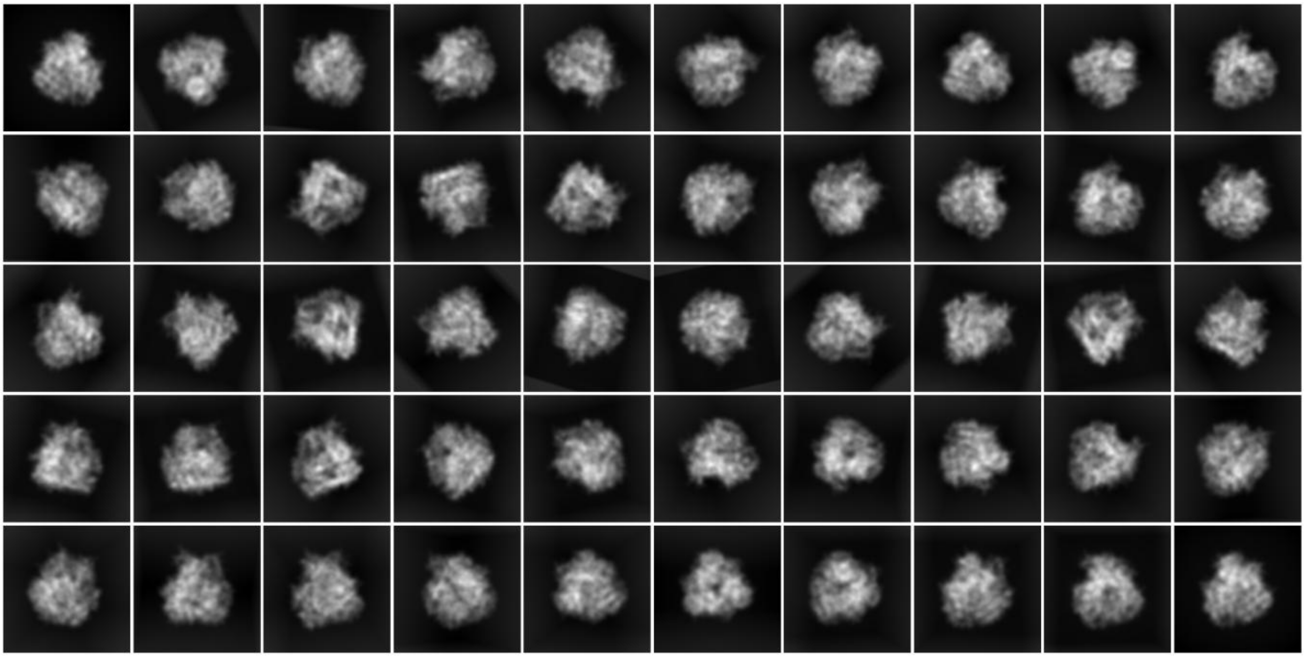

Created 2D templates from the designed model
